# Rapid formation of peptide/lipid co-aggregates by the amyloidogenic seminal peptide PAP_248-286_

**DOI:** 10.1101/2020.03.04.976746

**Authors:** E.W. Vane, S. He, L. Maibaum, A. Nath

**Affiliations:** University of Washington

## Abstract

Protein/lipid co-assembly is an understudied phenomenon that is important to the function of antimicrobial peptides as well as the pathological effects of amyloid. Here we study the co-assembly process of PAP_248-286_, a seminal peptide that displays both amyloid-forming and antimicrobial activity. PAP_248-286_ is a fragment of prostatic acid phosphatase and has been reported to form amyloid fibrils, known as semen-derived enhancer of viral infection (SEVI), that enhance the viral infectivity of HIV. We find that in addition to forming amyloid, PAP_248-286_ much more readily assembles with lipid vesicles into peptide/lipid co-aggregates that resemble amyloid fibrils in some important ways but are a distinct species. The formation of these co-aggregates, which we term “messicles”, is controlled by the peptide:lipid (P:L) ratio and by the lipid composition. The optimal P:L ratio is around 1:10 and at least 70% anionic lipid is required for co-aggregate formation. Once formed, messicles are not disrupted by subsequent changes in P:L ratio. We propose that messicles form through a polyvalent assembly mechanism, where a critical surface density of PAP_248-286_ on liposomes enables peptide-mediated particle bridging into larger species. Even at ~100-fold lower PAP_248-286_ concentrations, messicles form at least 10-fold faster than amyloid fibrils. It is therefore possible that, some or all of the biological activities assigned to SEVI, the amyloid form of PAP_248-286_, could instead be attributed to a PAP_248-286_/lipid co-aggregate. More broadly speaking, this work provides a potential framework for the discovery and characterization of peptide/lipid co-aggregates by other amyloid-forming proteins and antimicrobial peptides.

**Statement of Significance:** PAP_248-286_, a fragment of prostatic acid phosphatase, forms amyloid thought to enhances the infectivity of many viruses, including HIV. This amyloid, termed semen-derived enhancer of viral infection (SEVI), has been assigned responsibility for all of PAP_248-286_’s biological activities, while the monomer is thought to be inactive. However, SEVI formation is quite slow and requires very high concentrations of PAP_248-286_. Here, we show that PAP_248-286_ can instead assemble much more rapidly with lipid membranes to form another species, mechanistically and morphologically distinct from both monomer and SEVI amyloid. We have characterized this new species, which could play a role in the biological activities currently ascribed to SEVI. Additionally, our proposed mechanism for peptide/lipid co-assembly could apply to other biologically important systems.

## Introduction

Protein self-assembly has been important area of study for the past century. It is ubiquitous in biology – essential for normal function and yet also underlying major diseases. Structural filaments like actin and tubulin microtubules, whose assembly is critical for proper cellular motility, morphology, and transport and division, have been the topics of study for over 70 years (1, 2). Aggregation into amyloid filaments, a hallmark of disorders such as Alzheimer’s disease (AD), Parkinson’s disease (PD) and type II diabetes, has been studied for even longer (3). Self-assembly can serve to regulate metabolism, as the assembly of metabolic enzymes into fibers or foci in response to different environmental conditions can affect their activity (4). Recently, there has also been a resurgence of interest in the co-assembly of proteins with other macromolecules (often charged polymers) into phase-separated liquid droplets inside the cell. These membrane-less organelles are able to self-assemble and condense in a tunable, regulated fashion, so as to carry out specific metabolic or regulatory functions (5–7). Here, we focus on a related but distinct phenomenon, which is relevant both to amyloid pathology and to the function of antimicrobial peptides: the co-assembly of proteins and lipids into mixed aggregates.

There are as many as 40–50 amyloid-associated diseases, wherein diverse proteins misfold and self-assemble into characteristic insoluble, fibrillar, β-sheet-rich aggregates (8). The toxicity in these diseases is generally thought to result from the remodeling or disruption of biological membranes by pre-fibrillar or non-fibrillar oligomeric states rather than mature amyloid fibrils (9–11). Amyloid fibrils are relatively inert once formed, and can play important functional roles including in bacterial biofilm structure, peptide hormone storage, and protection against microbial infection (12–17).

Antimicrobial peptides (AMPs) are an important component of the innate immune system in animals ranging from insects to mammals, and are also produced by plants, fungi, and bacteria. Aside from immunomodulatory behavior, AMPs provide protection from pathogens by binding to and penetrating or disrupting pathogen membranes and/or by self-assembling around pathogen cells and causing them to agglutinate (15, 18–25). This agglutination disrupts the ability of pathogens to infect their hosts, increases the local peptide concentration, and makes them better targets for processes such as phagocytosis (15, 26, 27). Assemblies formed by AMPs that can operate via an agglutination mechanism, such as eosinophil cationic protein or LL-37, bear a striking resemblance to amyloid aggregates (21, 22). AMPs and amyloid-forming peptides share important physicochemical and functional features (16, 28). Indeed, the AD-associated peptide amyloid-β, displays potent agglutinative antimicrobial activity mediated by amyloid fibrils (15, 29). There is growing interest in the idea that some pathological amyloid-forming peptides confer protection against microbial pathogens, and that protein aggregation diseases may be the undesirable side effects of otherwise beneficial antimicrobial activity.

Peptide/lipid co-assembly impacts both amyloid pathology and AMP activity. Lipid membranes are known to promote aggregation of various amyloid-forming peptides (30, 31), and can alter the morphology of the resulting aggregates (32–34). In many cases, the lipid remains closely associated with the aggregated peptide (35, 36). Stable tau/lipid oligomers and α-synuclein lipopeptide nanoparticles have been suggested to be important in the pathogenesis of tauopathies and synucleinopathies, respectively (37–39). Agglutinative antimicrobial activity also requires the co-assembly of AMPs with bacterial membranes. However, little is understood about the physical processes that drive peptide/lipid co-assembly, or about the mechanism of the co-assembly formation.

Here, we study the phenomenon of peptide/lipid co-assembly using a peptide fragment of prostatic acid phosphatase (PAP_248-286_) as a model system. PAP_248-286_ is a peptide found in semen that displays antimicrobial activities via bacterial agglutination (40). PAP_248-286_ is also known to form amyloid. Amyloid formed by the PAP_248-286_ monomer, known as semen-derived enhancer of viral infection (SEVI), was discovered to be responsible for the large enhancement of HIV infectivity observed in seminal fluid (41). Since its discovery in 2008, SEVI has also been shown to enhance cytomegalovirus, herpes simplex virus type 1 and 2, and ebola virus infection as well as participate in sperm quality control (42–45). The exact mechanism for these activities is still unknown, but it has been hypothesized that the cationic SEVI amyloid can act like a bridge, trapping or bringing cells together (46). All of PAP_248-286_’s biological activity has been assigned to SEVI amyloid. However, the PAP_248-286_ monomer also enhances HIV infection after seminal fluid is doped in (41). Seminal fluid is a complex matrix that contains many different macromolecules including membrane-enclosed extracellular vesicles (EVs), proteins, lipids, and in the case of viral infection, viral particles (47–49). We therefore believe that it is possible that the peptide/lipid membrane co-aggregation of the PAP_248-286_ monomer with a component of seminal fluid could be an important, yet overlooked mechanism for PAP_248-286_’s activity.

We show that, in addition to forming amyloid, PAP_248-286_ is also able to assemble with lipids into large PAP_248-286_/lipid co-aggregates. Using liposomes as a model lipid membrane system and a variety of biophysical techniques including negative stain electron microscopy (EM), dynamic light scattering (DLS), circular dichroism (CD) spectroscopy, and fluorescence correlation spectroscopy (FCS), we explore the particular regime under which these co-aggregates form. These experiments reveal that PAP_248-286_/lipid co-aggregate formation is dependent on the PAP_248-286_:lipid ratio, as well as lipid membrane composition. Finally, we propose a formation mechanism by which polyvalent PAP_248-286_-mediated interactions drive co-aggregation at an optimal peptide:lipid ratio. The assembly process and the conditions that favor it may be important in explaining the diverse biological activities observed for PAP_248-286_.

## Methods

### Materials

1-palmitoyl-2-oleoyl-glycero-3-phosphocholine (POPC) and 1-palmitoyl-2-oleoyl-sn-glycero-3-phospho-(1’-rac-glycerol) (POPG) were purchased from Avanti Polar Lipids. Other reagents were purchased from Fisher Scientific and were ACS reagent grade or better.

### PAP 248-286 preparation

Synthetic PAP_248-286_ was purchased from Genscript and New England Peptides. Peptide was dissolved in either Tris buffer (20 mM Tris, 100 mM NaCl, pH 7.4) or filtered milliQ water, flash frozen, and stored at −80°C. The final concentration of PAP_248-286_ stock was determined by absorbance at 280 nm.

### Liposome preparation

LUVs were prepared with either 100% POPG, 7:3 POPG:POPC, 1:1 POPG:POPC, or 100% POPC (Avanti Polar Lipids). Stock lipid solutions were prepared at 19 mg/mL in chloroform and stored at −20°C for 6 months-1 year. The total phosphorous content of the lipid stocks was verified using the “Determination of Total Phosphorus” analytical procedure outlined by Avanti Polar Lipids, Inc. adapted for a Biotek Synergy HTX multi-mode plate reader.

Liposomes were prepared by measuring the appropriate amount of each lipid stock in chloroform and evaporating bulk chloroform with nitrogen gas to form lipid films. These films were left under vacuum for 1-14 hours and then hydrated with Tris buffer. The rehydrated lipid film was then subjected to 10 freeze-thaw cycles, followed by 23 passes through a mini-extruder (Avanti Polar lipids) with a 100 nm membrane. The final concentration of LUVs was determined using the total phosphorous assay, described earlier. Liposomes doped with DiOC_16_ (ThermoFisher Scientific) were prepared in the same manner, except DiOC_16_ in DMSO was added to the lipid stocks in chloroform at a molar ratio of 1:135 (dye:lipid). Liposomes filled with calcein (MP Biomedicals) were prepared in the same manner but hydrated in Tris buffer with 7 mM calcein and 0.3 mM EDTA.

### Microscope imaging

1:1, 1:10, and 1:25 PAP_248-286_:PG samples were prepared by incubating 10 μM PAP_248-286_ with 10, 100, or 250 μM POPG liposomes for 30 minutes. Samples were placed on a coverslip and imaged using a 20x objective of a home built, inverted Zeiss Axio Observer D1 confocal microscope equipped with an Andor iXon3 camera and Micro-manager open source microscopy software. Images were further processed with Fiji software.

For dissociation in the presence of excess PAP_248-286_, PAP_248-286_ was centrifuged at 21.1k x g for 30 minutes to remove any aggregates. “Dissociation” and “Dilution control” samples were prepared by incubating 20 μL of 20 μM messicles (20 μM PAP_248-286_ and 200 μM POPG liposomes in Tris buffer) for 4 minutes at room temperature. After incubation, 80 uL of 20 μM PAP_248-286_ was added to the dissociation sample, bringing the final concentration to 20 μM PAP_248-286_ and 40 μM POPG. 80 uL of tris buffer was added to the dilution control, bringing the final concentration to 4 μM PAP_248-286_ and 40 μM POPG. The “fresh 1:2” sample was prepared as a concentration match to the dissociation sample. It was prepared by incubating 100 μL of 20 μM PAP_248-286_ with 40 μM POPG for 4 minutes at room temperature. Dissociation in the-presence of excess POPG liposomes samples were prepared in a similar manner. Here, the dissociation sample was prepared by adding 80 μL of 200 μM POPG to the 20 uL of 20 μM Messicle, bringing the final concentration to 4 μM PAP_248-286_ and 200 μM POPG. The concentration match for the dissociation sample, called “fresh 1:50”, was prepared by incubating 100 μL of 4 μM PAP_248-286_ with 200 μM POPG for 4 minutes at room temperature. Samples were imaged 4, 30, and 60 minutes after the final solution was prepared. Data presented are representative of seven different images collected throughout different x,y-coordinates in the sample droplet.

### Electron microscopy

300 mesh carbon-coated copper grids (Electron Microscopy Services) were glow-discharged and spotted with 3 μL of buffer, 22 μM PAP_248-286_, 10 μM PAP_248-286_ with 10 μM POPG, 100 μM POPG, 100 μM 7:3 POPG:POPC, or 250 μM POPG, and 250 μM POPG liposomes after a 4 minute incubation, and SEVI amyloid diluted to 0.01 mg/mL. Samples adsorbed for 60 seconds, then were blotted by filter paper, rinsed with filtered milliQ water, blotted with filter paper, and then floated on a drop of NanoW stain (NanoProbes) for 60 seconds. Stain was blotted with filter paper, and grids were allowed to dry on filter paper then stored in a container at room temperature. Grids were visualized on a Morgagni electron microscope with a 100kV beam.

### Sedimentation assay

20 μM messicles (20 μM PAP_248-286_ and 200 μM POPG), 200 μM POPG liposomes, and 20 μM PAP_248-286_ were incubated at room temperature for 1 hour. Each solution was split in half and the samples were centrifuged at 9000g for 10 minutes. The supernatant for each sample was removed and the pellet was resuspended in Tris buffer. The concentration of lipid in the pellet and supernatant was determined by the total phosphorous assay. The fraction of the total inorganic phosphate (P_i_) for each sample was calculated by dividing the lipid concentration in either the pellet or supernatant by the sum of the two. Data presented are the average of 2 independent experiments.

Messicle disruption assays were conducted in a similar manner, except after the supernatant was separated from the pellet, the absorbance and fluorescence of the both fractions were measured. Supernatant and pellet were spotted onto a Take3 micro-volume plate (BioTek). The plate was placed into a BioTek Synergy HTX plate reader and the calcein absorbance spectra were collected from 450-550 nm. Calcein fluorescence was then measured using 460 nm and 528 nm filters for excitation and emission, respectively. Data presented are the average of 2 independent experiments.

### Dynamic light scattering

PAP_248-286_ was first centrifuged at 21.1k x g for 30 minutes to remove any large aggregates. DLS titrations of 10 μM PAP_248-286_ with 10, 50, 100, 150, or 200 μM of 100% POPG, 7:3 POPG:POPC, 1:1 POPG:POPC, or 100% POPC were prepared in Tris buffer and incubated for 4 minutes at room temperature. Samples were added into a 1 μL Quartz Cuvette (Wyatt Technology) and measurements were collected with a DynaPro NanoStar (Wyatt Technology). Ten to fifteen 10-second acquisitions were measured at 25°C and averaged. The diffusion coefficient (*D_t_*), in nm^2^/μs, of each sample was determined by re-fitting the average autocorrelation curve with a single component fit using GraphPad Prism:

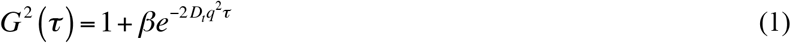

where *β* is the intercept and ***τ*** is the delay time. The scattering vector, *q*, is constant for the instrumental setup,

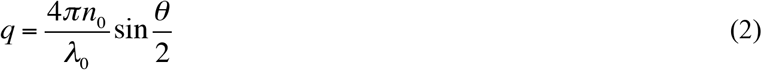

where the refractive index of the solution *n*_0_ is set to 1.33, the wavelength of the laser *λ*_0_ is set to 661 nm, and the scattering angle θ is set to 90°. The Stokes radius (*R_h_*) was then calculated using the Stokes-Einstein relation,

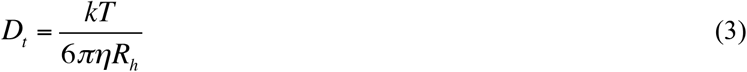

where *k* is Boltzmann’s constant, *T* is temperature, and the solution viscosity *η* is set to 0.89 mPa·s (50).

For dissociation in the presence of excess PAP_248-286_, PAP_248-286_ was centrifuged at 21.1k *g* for 30 minutes. “Dissociation”, “Dilution control”, and “Fresh 1:2” samples were prepared in the same manner as they were for the microscope dissociation experiment. Measurements were collected 4 minutes and 30 minutes after the final solution was prepared and in the same manner as previously stated. Analysis was performed as described above.

### Circular dichroism

CD samples of 10 μM PAP_248-286_ with 0, 50, 100, 150, 200, or 1000 μM of 100% POPG, 1:1 POPG:POPC, or 100% POPC liposomes were prepared in Tris buffer and incubated at room temperature for 4 minutes. Samples were measured in a 1 mm cuvette and spectra were collected on a Jasco J-1500 CD spectrometer. Measurements were taken at 25°C from 250-155 nm in continuous scanning mode with a scanning rate of 100 nm/min, data integration time of 1 second, and bandwidth of 1 nm. Data presented are an average of 5-7 scans after subtraction of the liposome background spectra for each sample.

### ThT kinetic assays

A 500 μM solution of Thioflavin T (ThT) was prepared in Tris buffer and filtered through a 0.22 μm syringe filter.

Tris buffer was added to wells of a 96-well black with clear, flat-bottom non-binding plate (Corning). PAP_248-286_ was centrifuged at 21.1k x g for 30 minutes and added to each well, followed by ThT and either 1:1 POPG:POPC or POPG liposomes, bringing the final volume to 200 μL. The final concentration of ThT, PAP_248-286_, and 1:1 POPG:POPC or POPG in each well was 25 μM, 10 μM, and 100 μM, respectively. Additional wells were prepared with ThT and 1:1 POPG:POPC or POPG as a lipid background. The plate was covered with clear polyolefin sealing tape (Thermo Scientific) to prevent evaporation and placed in a Biotek Synergy HTX multi-mode plate reader. Samples were held at 25°C and fluorescence was measured every 5 minutes using 440 nm and 485 nm filters for excitation and emission, respectively. Immediately before each read, the plate was shaken for 10-60 seconds. Since liposomes enhance ThT fluorescence, the lipid background control was first subtracted from each time point. The fluorescence signal at time 0 was then subtracted from the background subtracted values, yielding the ΔRFU over time. Replicates were plotted using GraphPad Prism.

For the SEVI fibril ThT kinetics, a 2.5 mM solution of ThT was prepared in PBS buffer (137 mM NaCl, 2.7 mM KCl, 10 mM Na_2_HPO_4_, 1.8 mM KH_2_PO_4_, pH adjusted to 7.4 with HCl), filtered through a 0.22 μm syringe filter, and stored at 4°C for up to 1 week. 1.1 mM PAP_248-286_ in PBS buffer was centrifuged at 21.1k x g for 30 minutes. 7% (v/v) of Tris buffer was added to bring the final volume to 107 uL and the final concentration of PAP_248-286_ to 1 mM. For ThT fluorescence measurements, 2.5 mM ThT was diluted to 25 μM ThT in PBS, filtered through a 0.22 μm syringe filter, and added to wells of 96-well black with clear, flat-bottom non-binding plate (Corning). As a ThT background, the fluorescence of this solution was measured using a 440 nm excitation and 485 nm emission filter on a Biotek Synergy HTX multi-mode plate reader. 2 uL of 1 mM PAP_248-286_ was then added to each well, bringing the final concentration 11 μM PAP_248-286_. The plate was placed in the Biotek plate reader and shaken linearly at 567 cpm for one minute and the ThT fluorescence in the presence of PAP_248-286_ was measured again. This process was repeated at each time point. Between time points, 1 mM PAP_248-286_ was agitated at 1200 RPM on an Eppendorf Thermomixer C at 37°C. For analysis, the background ThT fluorescence was subtracted from the ThT fluorescence in the presence of PAP_248-286_ and plotted using GraphPad Prism.

### Messicle seeding experiments

2.2 mM PAP_248-286_ was dissolved in 0.22 μm syringe filtered milliQ water and centrifuged at 21.1 kg for 30 minutes. After centrifugation, the 2.2 mM PAP_248-286_ was added to five separate tubes to be used for seeding experiments. (An equal volume of syringe filtered milliQ water was added to five separate tubes to serve as negative controls.) The concentration of the remaining solution was verified by its absorbance at 280 nm, and it was used for preparation of messicle and PAP_248-286_ stock solutions.

Seed, messicle, PAP_248-286_, POPG, and buffer control stock solutions were prepared at twice the desired concentrations. Stocks were prepared by adding 0.22 μm syringe filtered PBS, POPG liposomes in Tris, or Tris buffer, and PAP_248-286_. The solutions were allowed to incubate at room temperature while the seeds were prepared. Days prior to the seeding experiment, SEVI fibrils were prepared by shaking 1.1 mM PAP_248-286_ at 1200 RPM on an Eppendorf Thermomixer C at 37°C until the solution became cloudy. Fibril formation was confirmed by an increase in ThT fluorescence, and fibril samples were stored at 4°C. Seeds were prepared by vortexing SEVI fibrils for 4 minutes. The seeds were then added followed by the addition of 1 part of each stock solution to 1 part 2.2 mM PAP_248-286_ or PBS, bringing the final concentrations to 1.1 mM PAP_248-286_ and 5% Tris buffer.

For ThT fluorescence measurements, the 2.5 mM ThT solution was diluted to 25 μM ThT in PBS, filtered through a 0.22 μm syringe filter, and added to wells of 96-well black with clear, flat-bottom non-binding plate (Corning). As a ThT background, the fluorescence of this solution was measured using 440 nm excitation and 482 nm emission wavelengths on a Molecular Devices Spectramax Gemini EM plate reader. Each sample or control was then added the wells, bringing the final concentration PAP_248-286_ concentration to 11 μM. The plate was shaken linearly at 567 cpm for one minute and the ThT fluorescence was measured again. This process was repeated for each time point. Between time points, samples were agitated at 1200 RPM on an Eppendorf Thermomixer C at 37°C. For analysis, the background ThT fluorescence was subtracted from the ThT fluorescence in the presence of PAP_248-286_ and plotted using GraphPad Prism.

### Fluorescence correlation spectroscopy

Alexa Fluor 488 NHS Ester (Life Technologies) was conjugated to 100 μM PAP_248-286_ in a sodium phosphate buffer (20 mM sodium phosphate, 50 mM NaCl, pH 6.1) by incubating ~3-fold excess dye at room temperature for 3 hours or overnight at 4°C. The reaction was quenched with the addition of a pH 8.0 buffer containing 40 mM Tris and 50 mM NaCl. Labeled PAP_248-286_ was separated from unlabeled PAP_248-286_ and free dye on a BDS Hypersil C18 column run with a water/methanol gradient in the presence of 0.1%TFA on a Dionex Ultimate 3000 HPLC. Fractions containing labeled protein were collected, and solvent was evaporated with a Savant SC210A SpeedVac Concentrator with in-line Savant RVT5105 Refrigerated Vapor Trap. Aliquots of dried, labeled PAP_248-286_ was stored at −80°C until immediately prior to use.

FCS measurements were collected on a home-built instrument consisting of an inverted Zeiss Axio Observer D1 (Carl Zeiss Microscopy) microscope equipped with a Picoquant PDL 828 Sepia II pulsed laser driver, Hydra-Harp 400 detection electronics, and a Tau-SPAD photon counting detector (PicoQuant GmbH). The laser was passed through a 488 nm/10 nm excitation filter and collected by a 488 nm dichroic mirror and 535 nm/70 nm emission filter (all filters from Chroma Corp).

50 μL samples containing 2 μM unlabeled PAP_2_4_8_-_28_6, a fixed nanomolar concentration A488-labeled PAP_248-286_ resuspended in a 20 mM Tris, 100 mM NaCl, pH 7.4 buffer, and different concentrations 100% POPG, 7:3 POPG:POPC, and 1:1 POPG:POPC liposomes were prepared in a Tris buffer (20 mM Tris, 100 mM NaCl, pH 7.4) and incubated for two minutes at room temperature. 5-7 consecutive 30 second measurements were collected and autocorrelation and time-correlated single-photon counting of the fluorescence signal from the detector were performed using the HydraHarp 400 and SymphoTime 64 software (PicoQuant). Liposomes were fluorescently labeled by incubating them with Nile Red (Acros Organics) in MeOH at a 1:100 molar ratio (dye:lipid) ratio for at least half an hour. Nile Red-labeled liposomes were run in the same manner as the binding titration samples to determine the diffusion time of a liposome.

Autocorrelation curves were fit using GraphPad Prism to the following equation for free diffusion of two fluorescently labeled components:

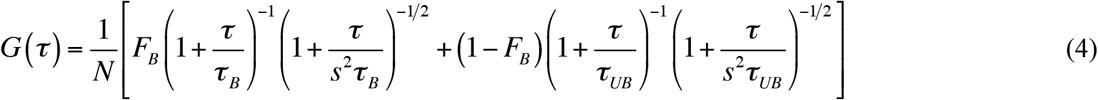

where *G*(*τ*) is the autocorrelation given lag time *τ, N* is average number of particles in the focal volume, *F_B_* and *τ_B_* are the fraction and diffusion time of liposome-bound PAP_248-286_, respectively, *τ_UB_* is the diffusion time of unbound PAP_248-286_, and *s* (the ratio of the radial to axial dimension of the focal volume) is fixed to 0.2 based on calibration measurements. *τ_B_* and *τ_UB_* were fixed, while *N* and *F_B_* were allowed to float. A one site binding model from GraphPad Prism was used to extract an apparent binding affinity, *K_D_*, from a plot of the fraction bound, *F_B_*, against the lipid concentration, [L]:

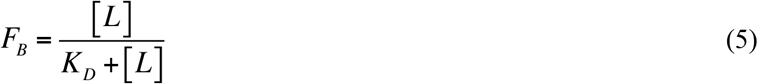

### Fluorescence burst analysis

Samples for dissociation in the presence of excess POPG liposomes were prepared in the same manner as they were for microscope imaging, except each sample contained nanomolar DiOC_16_-labeled lipids. Here, the dissociation and dilution control samples were prepared by adding unlabeled POPG or buffer to 20 μM Messicles containing DiOC_16_-doped POPG. Messicle, fresh 1:50, and lipid only controls were prepared as previously discussed, but contained an equivalent concentration of DiOC_16_-labeled lipids. After a 5 minute incubation of the final solution, the samples were placed onto the home-built Zeiss Axio Observer D1 microscope and 3 consecutive 10 minute fluorescence intensity time traces were recorded. Sample preparation and fluorescence intensity time trace collection was repeated twice for each sample, yielding an hour of time traces for each sample. A Python script was used to extract the sizes of fluorescence bursts from the intensity time traces above a defined threshold size. The size of each fluorescent burst for each sample was plotted as a frequency histogram using GraphPad Prism. A threshold of 200 photons/burst was selected because it yielded the best signal-to-noise ratio.

### Computer simulations

Molecular Dynamics computer simulations were performed using the Gromacs 5.1.4 software package (51) and employed the Charmm36 force field (52). Simulations were performed at a temperature of 303 K and at constant pressure and zero surface tension. The initial configuration of the phospholipid bilayer was constructed with the help of the Charmm-GUI web server (53). The bilayer contains 100 POPC and 100 POPG lipids, and was solvated in 150 mM KCl solution. The starting conformation of the peptide was an experimental structure (PDB entry 2L77 (54)), which was equilibrated under the same solvent conditions. The peptide was then placed above the phospholipid bilayer. Within approximately 50 ns the peptide absorbed onto the bilayer. The data analyzed in this work was obtained by averaging the trajectory from 100 ns up to 2000 ns of simulation time. We calculated the number of lipid molecules in contact with the peptide, defined as any lipid that has at least one atom within 2.5Å of any peptide atom.

## Results

### PAP_248-286_ rapidly forms large co-aggregates at specific peptide:lipid ratios

To understand its self-assembly in the presence of lipid membranes, PAP_248-286_ was incubated with 100 nm lipid vesicles with varying ratios of the anionic lipid POPG. A solution containing 10 μM PAP_248-286_ and 100 μM POPG (1:10 P:L ratio) became visibly cloudy within 15 minutes, while solutions at 1:1 or 1:25 P:L ratios remained clear (Figure 1a). Light microscopy showed that this cloudiness is due to the formation of micron-scale cloud-like or flocculent aggregates. These aggregates were not observed at the 1:1 P:L ratio, and are much less abundant at 1:25 than at 1:10 (Figure 1b). At 14,000x magnification, negative stain transmission electron microscopy (TEM) showed that aggregates at the 1:10 ratio look like masses of striated protein that appear somewhat fibrillar (Figure 1c). It should be noted that smaller aggregates or thinner, more ribbon-like aggregates were sometimes observed at a 1:1 and 1:25 P:L ratio, respectively. A large mass with a diameter of about 4 μm was also observed on the 1:10 P:L ratio grid, showing how heterogenous this species is (Figure 1d). This species is not formed by PAP_248-286_ or LUVs alone, showing that this aggregation is specific to peptide/lipid co-assembly (Figure S1). For convenience, we will refer to this mixed PAP_248-286_/lipid co-aggregated species as a “messicle” (an abbreviation for “*m*ultiple entrapped ve*sicles*” or for a “*mess* of ve*sicles*”).

**Figure 1:**
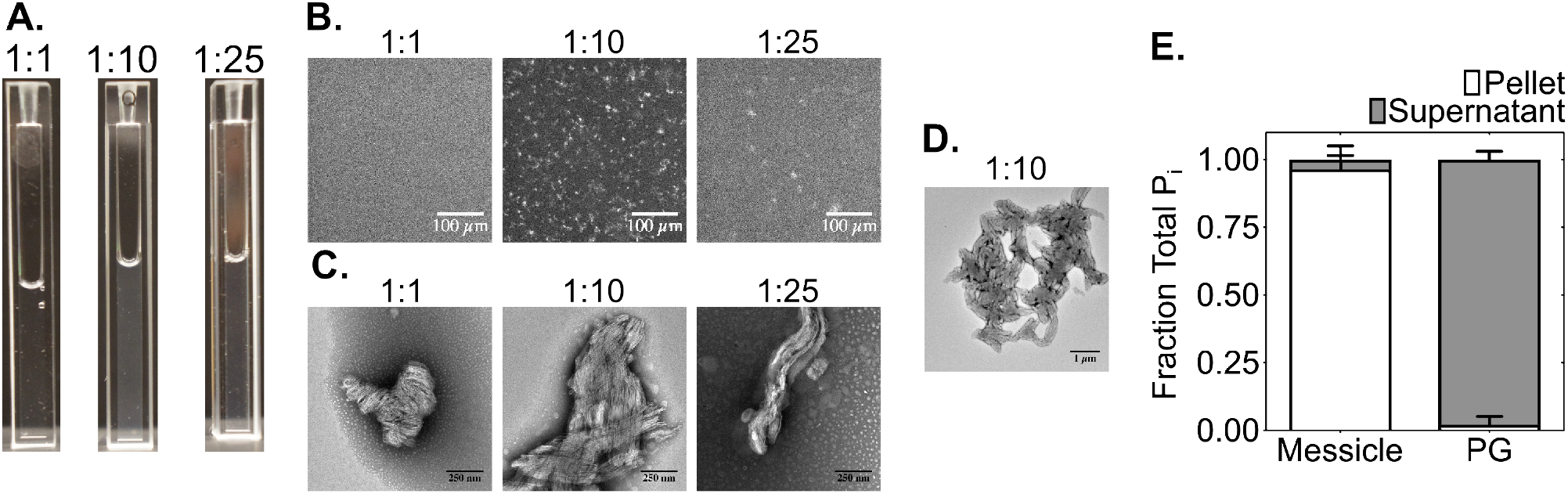
Large peptide-lipid co-aggregates, termed messicles, are preferentially formed at a 1:10 ratio of PAP248-286:POPG liposomes. (A) After a 15 minute incubation, 10 μM PAP248-286 mixed with 100 μM POPG liposomes is visibly cloudier than with 10 μM or 250 μM POPG. (B) After a 30 minute incubation, 20 μM PAP248-286 mixed with 200 μM POPG liposomes yields more numerous micron-scale aggregates than with 20 μM or 500 μM POPG. Images of the slide surface are at 20x magnification, and represent seven different fields of view taken at different positions on the slide surface. Scale bars indicate 100 μm. (C) Negative stain transmission electron microscopy (TEM) of 10 μM PAP248-286 incubated with 10, 100, and 250 μM POPG for 4 minutes shows that the 1:10 ratio also yields larger nm-scale aggregates. Scale bars indicate 250 nm. (D) TEM micrograph of a larger aggregate formed with 10 μM SEVI and 100 μM POPG, present on the same grid as the 1:10 image from panel C. Scale bar represents 1 μm. (E) Fraction of recovered phosphate (Pi) in pellet (white) or supernatant (gray) upon sedimentation of 20 μM messicle (20 μM PAP248-286 + 200 μM POPG) or 200 μM POPG liposomes after an hour-long incubation, indicating that the lipid has been almost entirely incorporated into the messicle co-aggregate. Bars represent the mean and standard deviation of two independent experiments.

To verify that messicles are indeed a peptide/lipid co-assembly, they were pelleted by centrifugation for 10 minutes at 9000g, and the concentration of phospholipids in solution vs. pelleted was determined using a total phosphorus assay. About 96% of phospholipid was pelleted with the messicles, while pelleted phospholipids were not detected in a control sample of POPG without PAP_248-286_ (Figure 1e). We also tested whether liposomes were disrupted upon incorporation into messicles, since PAP_248-286_ is known to disrupt lipid membranes at micromolar concentrations (55). POPG liposomes were filled with 7 mM of calcein, which results in partial self-quenching of this dye’s fluorescence. Messicles or control liposomes were again pelleted by centrifugation, the pellet was resuspended, and the calcein absorbance and fluorescence of the pellet and supernatant was measured. Calcein fluorescence was not detected in either pellet, but both supernatants showed similar calcein absorbance, suggesting that the liposomes in the messicles have leaked and released their calcein contents into solution. Fluorescence is higher for the messicle supernatant than the liposome control, indicating the liposomes in the control are intact and the encapsulated calcein remains partially quenched (Figure S2a and S2b). Beyond the observation that liposomes in vesicles have leaked their contents, the structure or arrangement of lipids in messicles remains unclear.

### Messicle formation depends on liposome charge

To explore the boundaries of messicle formation, we wanted to determine if this co-assembly forms with liposomes of other lipid compositions. A distinguishing characteristic of messicles is that they are large, which allowed the use of particle sizing by dynamic light scattering (DLS) to quickly test messicle formation under a variety of different lipid compositions and peptide:lipid ratios. In agreement with imaging at various magnifications (Figure 1), the largest particles in a titration of POPG into PAP_248-286_ form at the 1:10 P:L ratio. Moving away from that ratio yields smaller particles. Interestingly, messicle formation was not detectable when doing the same measurements with liposomes composed of 1:1 POPC:POPG, which carry a 50% negative charge, instead of the 100% negatively charged POPG liposomes. Messicle formation was also not observed with neutral POPC liposomes (Figure 2a). Therefore, we determined that, under our experimental conditions, messicles will not form with liposomes that have at or less than a 50% negative charge.

**Figure 2:**
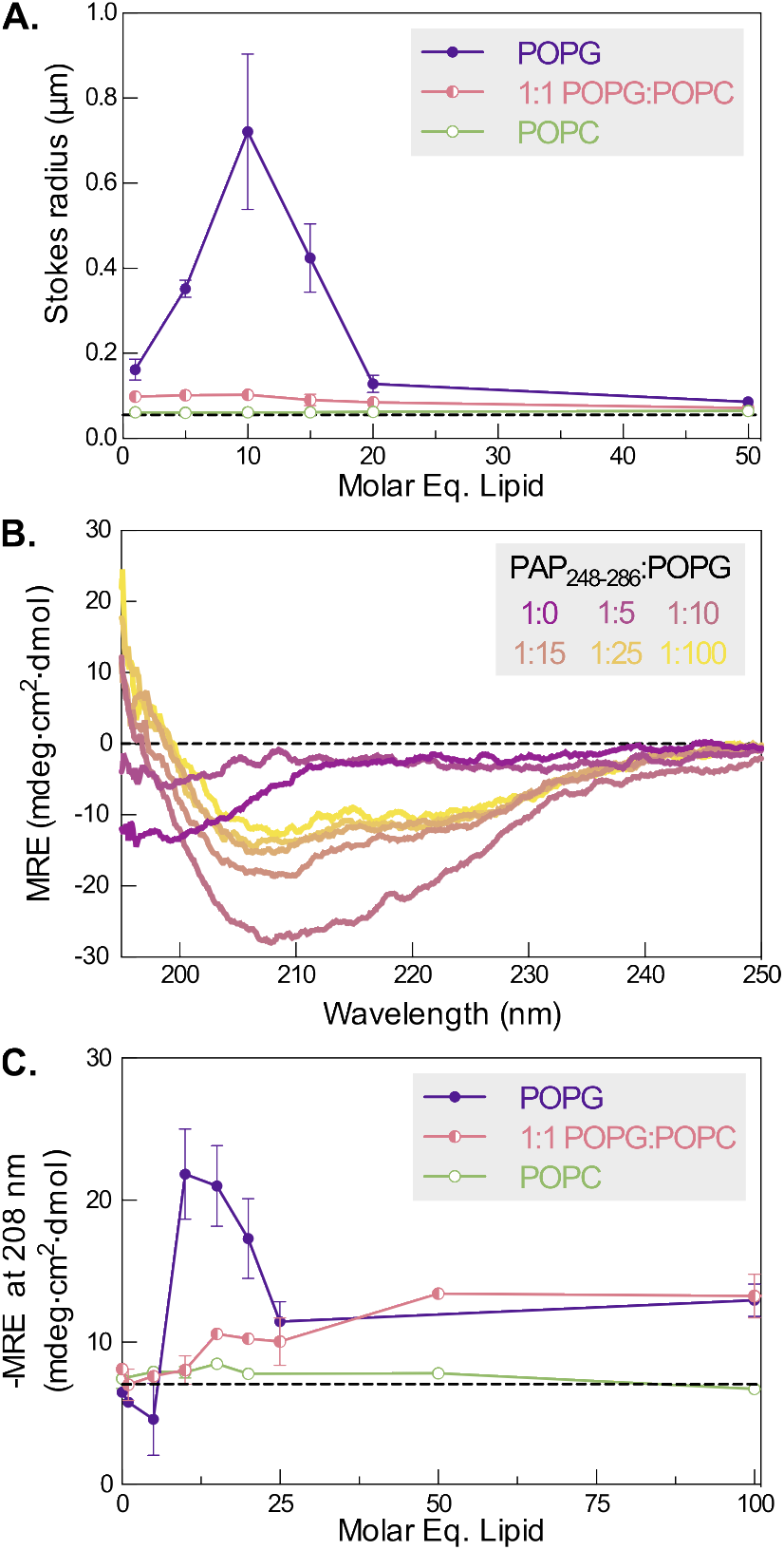
Messicle formation depends on the lipid charge. (A) Average particle size of 10 μM PAP_248-286_ incubated with 1, 5, 10, 15, 20, and 50 molar equivalents of POPG (*purple*), 1:1 POPG:POPC (*pink*), and POPC liposomes (*green*) for 4 minutes as determined by dynamic light scattering. Error bars represent the mean and standard deviation of two independent experiments for 1:1 POPG:POPC and POPC liposomes and four independent experiments for POPG liposomes. The black dashed line indicates the average Stokes radius measured for liposomes with 100 nm nominal diameter. (B) Representative overlay of far-UV circular dichroism (CD) spectra for 10 μM PAP_248-286_ incubated with varying concentrations of POPG liposomes for 4 minutes *(purple* to *yellow*). (C) The CD-derived negative mean residue ellipticity at 208 nm as a function of lipid concentration, for 10 μM PAP_248-286_ incubated for 4 minutes with POPG (*purple*), 1:1 POPG:POPC (*pink*), or POPC liposomes (*green*). Error bars represent the mean and standard deviation of two independent experiments for 1:1 POPG:POPC and POPC liposomes and four independent experiments for POPG liposomes. The black dashed line indicates the average – MRE at 208 nm for PAP_248-286_ alone.

PAP_248-286_ adopts a unique secondary structure in messicles. As monitored by far-UV circular dichroism (CD) spectroscopy, PAP_248-286_ displays a random coil spectrum in solution, and a classical α-helical spectrum at P:L ratios of 1:25 or greater. However, at the 1:10 P:L ratio with POPG, a large, distinctive signal with a minimum of 208 nm was observed after a 4-minute incubation (Figure 2b). The unusual spectral shape did not correspond to a single secondary structure, but could perhaps be due to a mixture of α-helix and β-sheet (56), β-sheet and random coil (57), or a combination of all three (58). The mean residue ellipticity at 208 nm was used to monitor the population of this secondary structure over different lipid compositions and P:L ratios (Figure 2c). A decrease in MRE at 208 nm is only observed with POPG and not with POPC or 1:1 POPC:POPG liposomes (Figure S3), again suggesting that messicles form near the 1:10 PAP_248-286_:lipid ratio but only for liposomes with > 50% negative charge.

### Messicles are amyloid-like, but distinct from bona fide SEVI amyloid

Because PAP_248-286_ forms amyloid fibrils (41) and anionic liposomes can promote amyloidogenesis of other proteins (30, 31, 33), we explored the possibility that messicles are, in fact, merely amyloid aggregates. Like most amyloid aggregates, messicles bind to and enhance the fluorescence of the diagnostic dye Thioflavin T (ThT) – in fact, to an even greater extent than PAP_248-286_-derived SEVI amyloid. This by itself does not establish that messicles are a variety of amyloid aggregate, since some non-amyloid states can also enhance ThT fluorescence (59, 60). Importantly, the formation kinetics are different: while PAP_248-286_ aggregation displays the characteristic sigmoidal formation kinetics when forming SEVI amyloid, messicle formation lacks a lag phase and instead displays exponential kinetics (Figure 3a). SEVI amyloid formation is relatively slow, taking ≥ 10 hours, even at ~1 mM peptide concentration under optimal conditions (when shaken at 37°C in phosphate buffered saline). By contrast, messicles form much more readily, taking ≤ 5 minutes at room temperature with just 10 μM peptide (in PBS, Tris or HEPES buffer). When visualized by negative stain EM, messicles were clearly more heterogenous and wider than the bundled SEVI fibrils, suggesting that messicles also have a distinct morphology from *bona fide* SEVI amyloid (Figure 3b).

**Figure 3:**
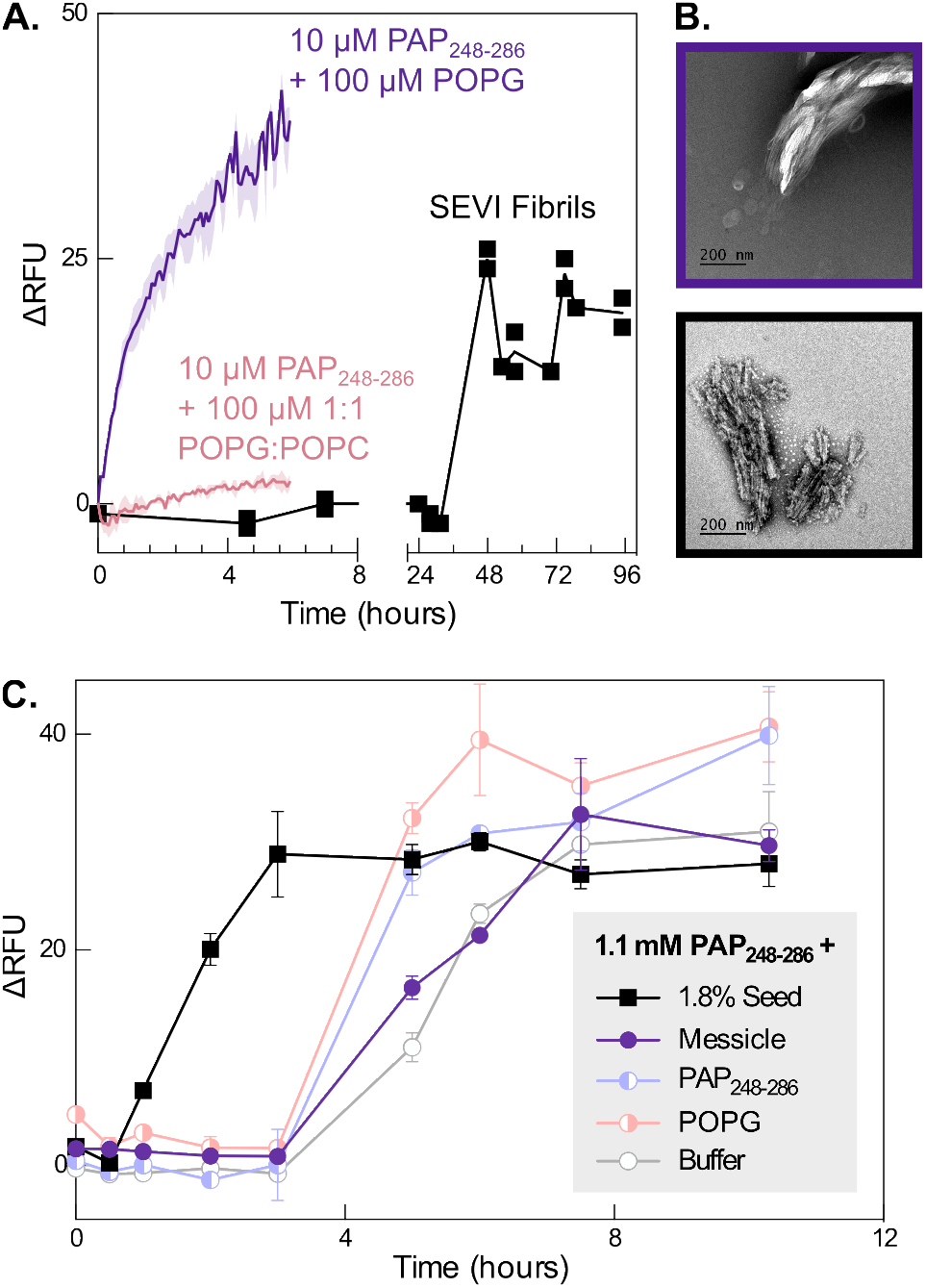
Messicles are kinetically and morphologically distinct from ***bona fide*** SEVI amyloid. (A) ThT fluorescence of 10 μM PAP_248-286_ with 100 μM POPG (messicle, *purple*) or 100 μM 1:1 POPG:POPC liposomes (*pink*), and 10 μM SEVI fibrils (*black*). The fluorescence signal was corrected for background from buffer, lipid and ThT. For the 10 μM PAP_248-286_ + 100 μM POPG or 1:1 POPG:POPC results, the error bands represent the mean and standard deviation of two independent experiments. The SEVI fibril trace is representative of four independent trials. (B) Representative negative stain EM micrographs of 10 μM messicles incubated for 1.5 hours (top, *purple*) and SEVI fibrils diluted to 2.1 μM (bottom, *black*), showing that messicles are more amorphous and less structured than amyloid fibrils. The magnification of both micrographs is 18,000x. (C) Representative seeding experiment containing 1.1 mM PAP_248-286_ with ThT and either 1.8% SEVI fibril seeds (20 μM PAP_248-286_, *black*), 20 μM messicle (20 μM PAP_248-286_ + 200 μM POPG, *purple*), 20 μM PAP_248-286_ (*blue*), 200 μM POPG (*pink*), or buffer (*grey*), showing that messicles are unable to seed amyloid formation as effectively as SEVI fibrils. The fluorescence signal was background corrected using analogous samples lacking the 1.1 mM PAP_248-286_. The error bars represent the standard deviation of three technical replicates.

Another characteristic feature of amyloid is that the addition of a small amount of aggregate seed to a solution of amyloid-competent protein monomers will nucleate the amyloid formation process, bypassing the lag phase in the sigmoidal kinetics (11). Regardless of the variability observed for SEVI amyloid kinetics (61), messicles were unable to seed SEVI amyloid formation to the same extent as an equivalent concentration of SEVI seeds. As controls, the two components of messicles, PAP_248-286_ and POPG, were also unable to seed amyloid formation (Figure 3c). This further suggests that messicles have a distinct molecular conformation from *bona fide* SEVI amyloid. Although they are ThT-positive, messicles are mechanistically and structurally distinct from amyloid fibrils.

### Messicle formation is not driven by charge neutralization

Given its strong dependence on liposome charge, one possibility is that messicle formation could be occurring through a charge neutralization mechanism, as seen in some flocculation phenomena (62). According to this mechanism, messicles from at the optimal PAP_248-286_:POPG ratio of 1:10 because the cationic peptide molecules exactly neutralize the negative charges on the outer leaflet of each liposome. Electrostatic repulsion between cationic particles at P:L ratios > 1:10, or between anionic particles at P:L ratios < 1:10, would disfavor condensation at other ratios.

This model predicts that messicles formed with 7:3 POPG:POPC liposomes would require ~1.4-fold less PAP_248-286_ to neutralize charge, since the surface charge is 30% less negative. In fact, the optimal ratio for messicle formation, assessed by DLS in six independent trials, was essentially identical whether using POPG or 7:3 POPG:POPC liposomes. (Figure 4a). While the precise optimal P:L ratio varied between 1:10 and 1:14 from day to day (presumably due to minor inconsistencies in liposome preparations), in every case the fold-change in the optimal ratio was much closer to 1 than the 1.4 predicted by the charge neutralization mechanism (Figure 4b). Messicles produced using POPG or 7:3 POPG:POPC liposomes are thus identical in both P:L ratio and morphology (Figure 4c). Because the optimal P:L ratio does not change with a changing liposome charge, a charge neutralization model is inconsistent with the data.

**Figure 4:**
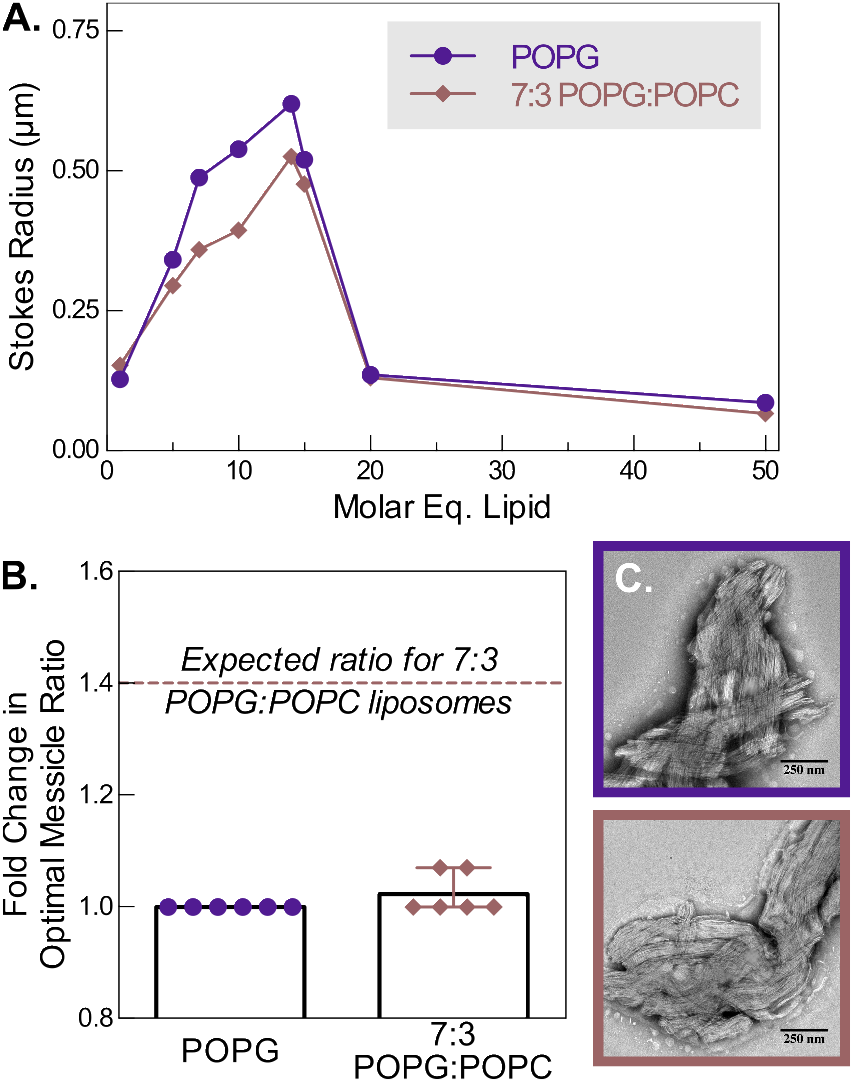
Co-assembly is not driven simply by charge neutralization-induced coagulation. (A) Representative particle size of 10 μM PAP_248-286_ incubated with 1, 5, 7, 10, 14, 15, 20, and 50 molar equivalents of POPG *(purple*) or 7:3 POPG:POPC (*brown*) liposomes for 4 minutes as determined by dynamic light scattering, showing that both lipid compositions have similar optimal ratios for messicle formation. (B) Across six independent experiments with either 10 or 20 μM PAP_248-286_, the optimal P:L ratio for messicle formation for 7:3 POPG:POPC is close to that for POPG, and never approaches the theoretical fold-change predicted by a charge neutralization model (dashed brown line). (C) Representative negative stain EM micrographs of 10 μM PAP_248-286_ incubated with 100 μM of POPG liposomes for 4 minutes (top, *purple*) and 10 μM PAP_248-286_ incubated with 100 μM of 7:3 POPG:POPC liposomes for 4 minutes (bottom, *brown*), showing that both messicle preparations are similar in size and morphology.

### Messicle formation is consistent with a polyvalent assembly model

Another general coagulation mechanism is interparticle bridging wherein a flocculant mediates the condensation of larger particles through multiple weak interactions (62). An optimal P:L ratio between 1:10 and 1: 14 corresponds to each PAP_248-286_ molecule making contacts with 5–7 lipid molecules (since the outer leaflet of a liposome contains roughly half of the total lipid). Molecular dynamics simulations of a single PAP_248-286_ molecule bound to a lipid bilayer (Figure 5a) show between 4 and 14 lipid contacts, meaning that a lipid contact number in the 5–7 range is consistent with the formation of a peptide monolayer surrounding each vesicle. Under these conditions, peptide-peptide interactions could bridge different liposomes, with multiple weak peptide-peptide interactions leading to condensation. An increase in lipid content would dilute the membrane-bound peptide so that the monolayers are incomplete, and make these bridging interactions less likely. Excess peptide would bind to PAP_248-286_-coated liposomes and passivate them, competitively blocking interactions with other peptide-coated liposomes and slowing down messicle formation (Figure 5b).

**Figure 5:**
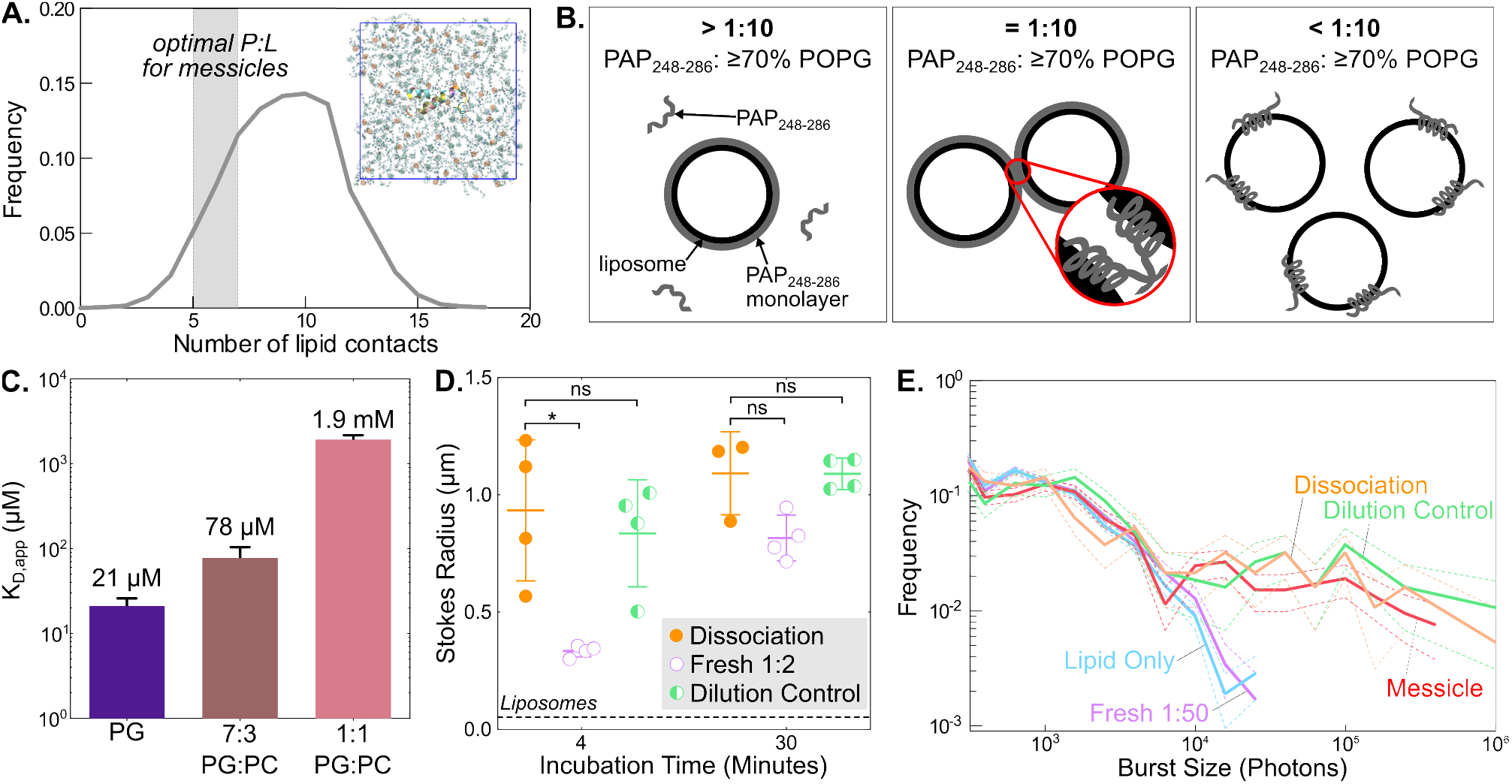
Messicle stability is consistent with PAP_248-286_-induced polyvalent assembly of liposomes model. (A) Probability distribution of observed peptide-lipid contacts in a simulation of a single PAP_248-286_ peptide absorbed onto a 50:50 POPC/POPG bilayer spans the range of optimal ratios for messicle formation, indicating that the conditions that drive messicle assembly are consistent with the formation of a peptide monolayer around each liposome. The inset shows a simulation snapshot, with a top-down view of the peptide and the upper bilayer leaflet. Lipids’ phosphate atoms are highlighted in orange (POPC) and green (POPG). (B) Schematic of the proposed mechanism. When the PAP_248-286_:lipid ratio is greater than 1:10 (left panel), there are too few liposomes to assemble or the excess PAP_248-286_ acts to passivate liposomes and slow down assembly. Near the optimal 1:10 PAP_248-286_:lipid ratio (center panel), if the anionic content is sufficiently high (≥ 70% POPG), the liposome surface is covered in a monolayer of peptide, allowing for many weak lipid-peptide-peptide-lipid interactions. When the PAP_248-286_: lipid ratio is less than 1:10, excess liposomes bind to and sequester PAP_248-286_. (C) The binding affinity of PAP_248-286_ for liposomes composed of POPG (*purple*), 7:3 POPG:POPC (*brown*), and 1:1 POPG:POPC (*pink*) determined by fluorescence correlation spectroscopy (FCS) lipid titrations (see Supplementary Figure 4). Each bar represents the mean K_D,app_ and SEM of at least 3 independent titrations. (D) Messicle stability upon the addition of excess PAP_248-286_ determined by DLS particle sizing. The sizes of the dissociated messicle, “dissociation”, (20 μM PAP_248-286_ + 40 μM POPG, *orange*), the concentration-matched control, “Fresh 1:2”, (20 μM PAP_248-286_ + 40 μM POPG, *purple*), and “Dilution control” (4 μM PAP_248-286_ + 40 μM POPG, *green*) were all measured after a 4 and 30 minute incubation period. The size of the POPG liposome is shown by the broken line. Each bar represents the mean and standard deviation of at least 3 independent replicates. ns, no statistically significant difference; *P <0.05 (Welch’s t test). (E) Messicle stability upon the addition of excess POPG determined by single particle fluorescence. Each sample contained 15 nM DIOC_16_-doped POPG liposomes. Burst size analysis was done for the 20 μM “Messicle” (20 μM PAP_248-286_ + 200 μM POPG, *red*), the dissociated messicle, “Dissociation”, (4 μM PAP_248-286_ + 200 μM POPG, *orange*), the lipid only control (200 μM POPG, *blue*), the “Dilution control” (4 μM PAP_248-286_ + 40 μM POPG, *green*), and the concentration-matched control, “Fresh 1:50”, (4 μM PAP_248-286_ + 200 μM POPG,*purple*). The lines represent the mean and standard deviation of 2 independent replicates collected over half an hour.

In this model, PAP_248-286_ needs to have a high enough binding affinity for the liposome to form a monolayer under the experimental concentrations. Fluorescence correlation spectroscopy (FCS) was used to assess liposome binding by measuring the change in diffusion time of fluorescently labeled PAP_248-286_ with liposomes of varying concentrations and compositions (Figure S5). The apparent K_D_ values of PAP_248-286_ for POPG, 7:3 POPG:POPC and 1:1 POPG:POPC were determined to be 0.02, 0.08 and 2 mM, respectively (Figure 5c). Binding was not observed with POPC liposomes (data not shown). The large difference in binding affinity of PAP_248-286_ for 7:3 and 1:1 liposomes could explain why messicles form with the former and not the latter; in our hypothesized model, the weaker affinity of PAP_248-286_ for 1:1 liposomes would disfavor monolayer formation, and therefore messicle assembly, under experimental conditions.

Under a polyvalent assembly model, with many weak interactions driving co-aggregation, messicles would dissociate slowly relative to the timescale of a single peptide dissociating from the membrane. To monitor messicle disassembly, we used DLS to measure particle size after the addition of excess PAP_248-286_ to pre-formed messicles, to bring the final P:L ratio to 1:2. If these messicles dissociated, their hydrodynamic radius would match that of freshly prepared 1:2 P:L; if not, hydrodynamic radius would match that of a “dilution control” to which a matched amount of buffer has been added. The hydrodynamic radius of the “dissociated” samples matched the dilution controls at both 4 minutes and 30 minutes, indicating that messicle dissociation is negligible on this timescale (Figure 5d). Note that the freshly prepared 1:2 P:L ratio sample continues to grow over 30 minutes, approaching the size of the other samples. This is in agreement with the observation that some messicle assembly can be observed at large P:L ratios (Figures 1c and 2a), albeit more slowly than at the optimal 1:10 ratio. These results are also in agreement with light microscopy (Figure S6a), where over the course of an hour, the dissociated species looks similar to the dilution control but the freshly prepared 1:2 sample continues to grow.

Unfortunately, DLS could not be used to test if addition of excess liposomes can cause messicle dissociation, because of interference from light scattered by the added liposomes. Instead, we applied confocal single-particle fluorescence burst analysis using fluorescently labeled liposomes. In this experiment, larger particles like messicles, containing multiple labeled liposomes, will yield more intense fluorescence bursts than smaller particles such as individual liposomes. We monitored messicles after the addition of POPG to bring the final P:L ratio of 1:50, and compared them to a freshly prepared 1:50 sample and to a “dilution control” of messicles to which buffer volume had been matched. Comparing the distribution of burst sizes collected over 30 minutes, the messicle sample resembles the dilution control, while the 1:50 sample resembles POPG liposomes alone (Figures 5e and S7). This result is also in agreement with light microscopy of each species over the course of an hour at a 20x magnification (Figure S6b). These results again indicate that once messicles have formed, they are resistant to disruption by excess peptide or lipid. This kinetic stability is consistent with a polyvalent interaction model.

## Discussion

We have discovered a new higher-order species of PAP_248-286_ that, while similar in some ways to the well-studied SEVI amyloid, has important structural and mechanistic differences. While PAP_248-286_ messicles are ThT positive, they are more heterogeneous than SEVI fibrils, and are unable to seed amyloid formation. They are also able to form at much lower concentrations (10 μM vs. 1.1 mM), more rapidly (< 30 minutes vs. ~8–48 hours), and with non-sigmoidal kinetics. Moreover, they have a distinct secondary structure. Under our conditions, we determined that messicles form most quickly near a 1:10 P:L ratio and require vesicles with at least 70% anionic lipids. However, messicles are relatively stable to subsequent changes in P:L ratio, consistent with a mechanism involving polyvalent assembly or interparticle bridging of lipid vesicles by PAP_248-286_.

We believe it is possible that some or all of the biological activity assigned to PAP_248-286_–derived SEVI amyloid is in fact due to messicle-like species. Semen enhances ThT fluorescence, an effect thought to indicate the presence of endogenous seminal amyloids including SEVI (63). However, it is also possible that some ThT fluorescence is due to the formation of amyloid-like messicles, since the *in vitro* ThT fluorescence of messicles is even stronger than that of SEVI amyloid (Figure 3A). Therefore, messicles may be involved in the most well-studied activity of PAP_248-286_: its ability to enhance HIV infection. *In vitro,* SEVI amyloid enhances HIV infection while freshly dissolved PAP_248-286_ monomer was inactive. The ability to enhance HIV infection was therefore assigned to SEVI amyloid and not monomeric PAP_248-286_. However, in the presence of seminal fluid, PAP_248-286_ also enhances HIV infection (41). It is therefore possible that enhancement of HIV infectivity is due to the co-assembly of PAP_248-286_ into a messicle or messicle-like species with a component of seminal fluid. Intriguingly, seminal fluid is abundant in extracellular vesicles (EVs), historically called prostasomes, which are negatively charged vesicles of diameter 40–500 nm, and could potentially serve as a component for messicle formation *in vivo* (47, 64, 65).

Similarly, the beneficial activities proposed for PAP_248-286_ could also depend on lipid-mediated co-assembly. It has been proposed that SEVI could be an antimicrobial peptide, and may participate in sperm quality control and removal. Like other self-assembling AMPs, SEVI fibrils can agglutinate bacteria and promote their phagocytosis (40). Via a similar mechanism, it was demonstrated that SEVI fibrils trap spermatozoa and increase their uptake by macrophages (45). However, the formation of messicle-like species presents a mechanism by which monomeric PAP_248-286_ could exert similar biological activities. Given our *in vitro* results, an agglutination mechanism driven by co-assembly would result in an atypical, highly biphasic dose-response curve for monomeric PAP_248-286_. Experiments carried out at a single concentration of monomeric PAP_248-286_ would fail to detect such activity unless both the peptide concentration and the ratio of peptide to target cells or vesicles happened to fall close to optimal values. Future work testing different peptide concentrations is required to reveal whether the PAP_248-286_ monomer can co-assemble with spermatozoa, pathogenic bacteria, or other constituents of seminal fluid to form messicle-like species.

It is also possible that many other factors could affect how easily and quickly messicles form. While we have determined a relatively narrow set of conditions under which messicles form, it is possible that they can also form in more complex systems. We have determined that both P:L ratio and lipid charge are important to the polyvalent co-assembly model that we have proposed. It remains to be seen if lipid compositions enriched in EVs, spermatozoa, or bacterial cell membranes affects messicle formation. In agreement with the work from Ramamoorthy and co-workers (55, 66), our results have demonstrated the importance of electrostatic interactions to the interaction of PAP_248-286_ monomer with membranes. It is possible that the negative charges associated with acidic sugars would also be able to drive co-assembly with PAP_248-286_ to form messicles. Gram-positive and -negative bacterial membranes, for example, are highly decorated in techoic acid polymers and lipopolysaccharide (LPS), respectively. It is also not known whether biologically relevant lipids such as sphingomyelins, phosphatidylinositols, and cardiolipin would impact PAP_248-286_/membrane interactions, and therefore the nature of PAP_248-286_/lipid co-assemblies.

While we have hypothesized that formation of a monolayer of peptide around liposomes results in interparticle bridging of these coated liposomes, we do not know the exact nature of the co-assembly. Another possible mechanism is one where peptides form a multimeric (perhaps amyloid-like) structure on the lipid membrane and these multimers are able to mediate interactions with other peptide-bound liposomes. It is also possible that these messicles are, at least in part, composed of dramatically restructured liposomes. We know that the liposomes within messicles are disrupted (Figure S2) and that other amyloid-forming peptides, like α-synuclein, are able to restructure lipid membranes (38, 39). Regardless of which of the proposed mechanisms is responsible for messicle formation, we would expect that polyvalent assembly is required. The relative stability of messicles to changes in P:L ratio (Figure 5) provides evidence for a polyvalent mechanism of co-assembly. It is possible that this is formation mechanism is generally applicable to other cases, as there are many other antimicrobial and amyloid-forming peptides that are able to bind to membranes and self-assemble. Therefore it is possible that many of the other biological functions and pathologies associated with these peptides could in fact involve messicle formation.

## Author Contributions

E.W.V., A.N. and L.M. designed research; E.W.V. and S.H. performed research; E.W.V., S.H., L.M. and A.N. analyzed data; E.W.V., A.N. and L.M. wrote the manuscript.

## Acknowledgements

The authors thank Drs. Hannah Baughman, Eri Nakatani-Webster, David Baggett and Shana Elbaum-Garfinkle for helpful comments, as well as Alexandra Walls and Drs. David Veesler and Justin Kollman for assistance with electron microscopy. The authors gratefully acknowledge support from an NSF Graduate Research Fellowship to E.W.V.

